# APC2 is Critical for Ovarian WNT Signalling Control, Fertility and Tumour Suppression

**DOI:** 10.1101/516286

**Authors:** Noha-Ehssan Mohamed, Trevor Hay, Karen R. Reed, Matthew J Smalley, Alan R. Clarke

## Abstract

Canonical WNT signalling plays a critical role in the regulation of ovarian development; mis-regulation of this key pathway in the adult ovary is associated with subfertility and tumourigenesis. The roles of Adenomatous polyposis coli 2 (APC2), a little-studied WNT signalling pathway regulator, in ovarian homeostasis, fertility and tumourigenesis have not previously been explored. Here, we demonstrate for the first time using constitutive APC2-knockout (*Apc2^−/−^*) mice, essential roles of APC2 in regulating ovarian WNT signalling and ovarian homeostasis. APC2-deficiency resulted in activation of ovarian WNT signalling and sub-fertility driven by intra-ovarian defects. Follicular growth was perturbed, resulting in a reduced rate of ovulation and corpora lutea formation, which could not be rescued by administration of gonadotrophins. Defects in steroidogenesis and follicular vascularity contributed to the subfertility phenotype. Tumour incidence was assessed in aged APC2-deficient mice, which also carried a hypomorphic *Apc* allele. APC2-deficiency in these mice resulted in predisposition to granulosa cell tumour (GCT) formation, accompanied by acute tumour-associated WNT-signalling activation and a histologic pattern and molecular signature seen in human adult GCTs. Hence, APC2 has an important tumour-suppressor activity within ovarian granulosa cells, most likely due to its role in regulating WNT-signalling. Importantly, given that the APC2-deficient tumours recapitulate the molecular signature and histological features of human adult GCTs, this APC2-deficient mouse has excellent potential as a pre-clinical model to study ovarian tumour biology and for therapeutic testing.

## Introduction

The canonical WNT signalling pathway is central to numerous biological processes and diseases (1). Within the ovary, it has been shown to be essential for female sex differentiation during embryogenesis (2–9), however, in the adult ovary its role is less well defined. Conditional deletion of β-catenin within murine granulosa cells of antral follicles did not affect folliculogenesis or ovulation (10, 11), but its removal within oviducts and uteri led to abnormalities therein, with lack of implantation sites rendering mice infertile as a result (11). Conditional deletion of *Wnt4* in ovarian granulosa cells or germline deletion of *Fzd4* in mice caused sub-fertility or complete infertility respectively (12, 13), but WNT signalling activity was not measured and it is unclear whether the reported phenotypes were caused by impaired ovarian canonical WNT signalling or by other mechanisms, potentially including non-canonical pathways. In mice with germline deletion of the WNT signalling agonist *Fzd1*, where 17.6% of female mice were infertile and characterized by early follicle depletion, but with no concomitant change in total activated β-catenin levels (14). Over-activation of canonical WNT signalling also has deleterious effects on ovarian homeostasis. Ovarian amplification of *Rspo1* (15), deletion of *Wnt5a* (antagonist of canonical WNT signalling) (16) or expression of dominant stable β-catenin (10, 17) all resulted in up-regulated ovarian WNT signalling and ovarian subfertility caused by disruption of follicle growth (16, 17), ovulation and luteinisation (10, 15). Taken together, these findings indicate the importance of tight regulation of canonical WNT signalling in growing follicles.

Human ovarian tumours are classified into epithelial ovarian cancers (90%), sex cord-stromal tumours (7%) and germ cell tumours (3%). Granulosa cell tumours (GCTs), which originate from granulosa cells of ovarian follicles, account for more than half of sex cord-stromal tumours (18). WNT signalling mis-regulation has been implicated in adult GCT formation, as several studies have demonstrated increased β-catenin protein levels therein, with nuclear localisation in some cases (17, 19, 20). A recent molecular study of GCTs showed epigenetic silencing of *DKK3*, the gene coding for the WNT-signalling antagonist Dickkopf, implying a need for WNT signalling activation in GCT development (25, 26). Furthermore, GEMMs in which WNT signalling was activated via the introduction of a gain of function mutation of R-spondin1 (15), or a degradation-resistant β-catenin (17), resulted in 15.8% or 57% of mice developing adult GCTs respectively.

Here, for the first time, we address the importance of APC2 in ovarian folliculogenesis, fertility and GCT formation. The ability of APC2 to regulate the β-catenin/WNT signalling pathway has been demonstrated in *Drosophila* and in cancer cell lines (21–25). Structurally, APC2 possesses AXIN1 and β-catenin binding sites, which enable it to destabilize β-catenin, targeting it for degradation and suppressing its transcriptional activity (26, 27), in addition to the APC-basic domain which enables it to regulate cytoskeleton and microtubule association (28–32) and spindle anchoring during mitosis (33). Importantly, however, in an *in-vivo* setting, APC2-dependent regulation of WNT signalling is tissue-specific, occurring in the liver and intestine but not in the mammary gland (34, 35). Little is known about how APC2 functions in adult ovaries, but APC2 loss has been reported in epithelial ovarian cancer (29, 36). Here, we show that *Apc2*-knockout mice (37) have a subfertility phenotype associated with an activation of ovarian WNT signalling, and that, on a hypomorphic *Apc* background (38, 39), loss of APC2 increases the incidence of ovarian GCTs which recapitulate the histologic pattern and molecular signature of human adult GCTs. Not only does this study extend our understanding of the tissue-specific regulation of WNT signalling, but also the APC2-deficient mouse has excellent potential as a pre-clinical model to study ovarian tumour biology and for therapeutic testing.

## Results

### APC2 deficiency results in sub-fertility

To evaluate the role of APC2 in the biology of the adult ovary, the impact of APC2 deficiency on normal ovarian homeostasis and fertility was first assessed. A retrospective analysis of mating efficiencies of wild type, heterozygous or homozygous breeding trios *(Apc2^+/+^, Apc2^+/-^* or *Apc2^−/−^* respectively) demonstrated that time between pairing mice and first litter production was significantly longer in *Apc2^−/−^* mice (Figure 1a). The number of gestations over the 3-month period following pairing was significantly reduced in *Apc2^−/−^* mice, with heterozygous mice also showing a reduction which did not reach significance (Figure 1b). Overall, there was a 40% reduction in the cumulative number of pups weaned over 3 months from *Apc2^+/−^* trios, compared to *Apc2^+/+^*, and this reduction was even more pronounced in *Apc2^−/−^* mice (Figure 1c). Indeed, one *Apc2^−/−^* trio was completely infertile over this period.

**Figure 1:**
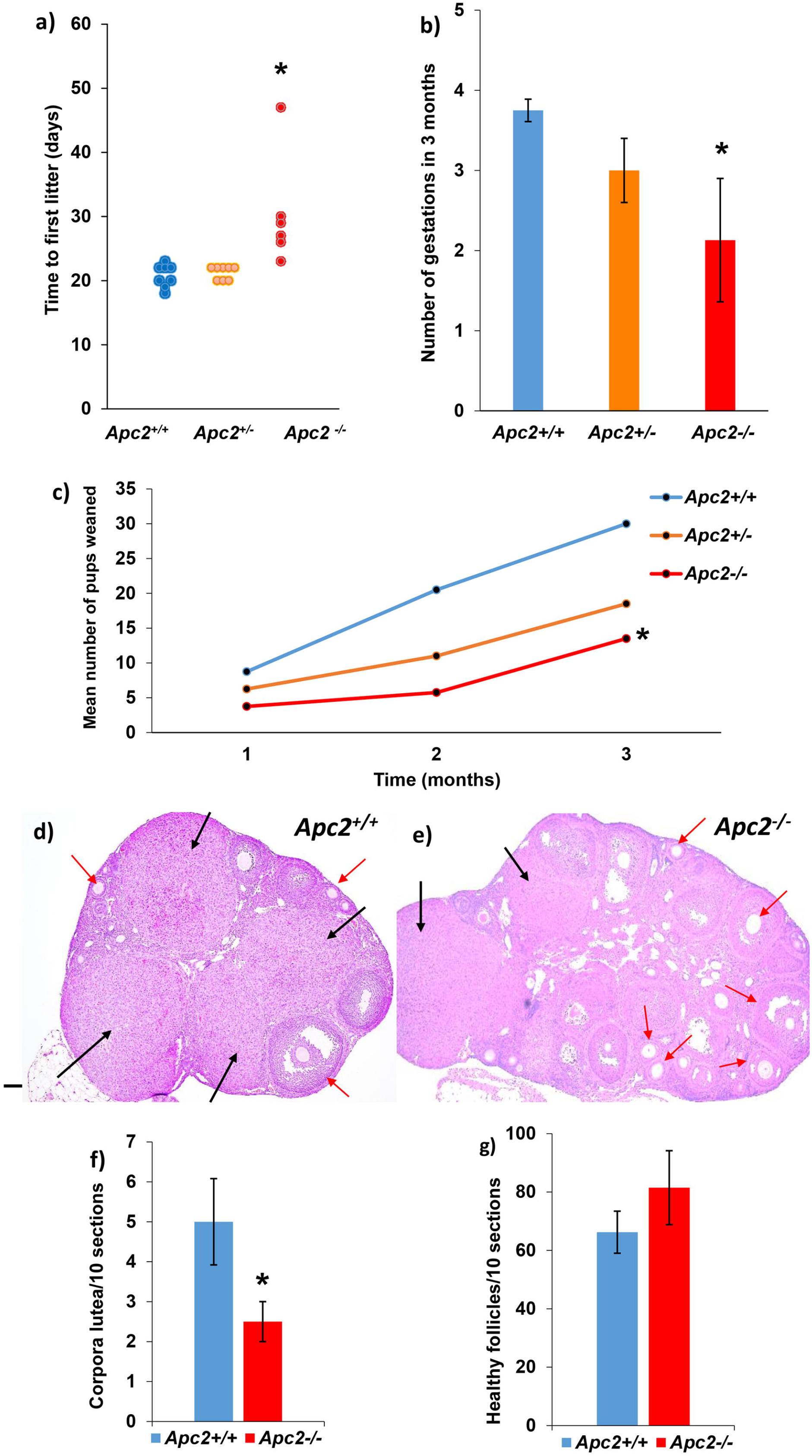
APC2 loss causes subfertility in adult female mice. **(a)** Mating efficiency of *Apc2* experimental genotypes as a function of time recorded in days between pairing the mice and delivering the first litter. N=4 breeding cages each with one *Apc2^+/+^, Apc2^+/−^* or *Apc2^−/−^* male crossed with 2 *Apc2^+/+^*, 2 *Apc2^+/−^* or 2 *Apc2^−/−^* female respectively. One *Apc2^−/−^* trio was completely sterile (*P<0.05). **(b)** Breeding efficiency as reflected by number of gestations occurring in a 3 month period (mean ± S.E, n=4). **(c)** Cumulative number of pups weaned in a 3 month period from 4 breeding pairs. Statistical significance between groups was determined using ANOVA test followed by Games-Howell post hoc analysis (variances of experimental groups were not homogeneous, tested by Levene’s test). **(d, e)** Representative photomicrographs of **(d)** *Apc2^+/+^* and **(e)** *Apc2^−/−^* ovaries, showing growing follicles (red arrows) and corpora lutea (black arrows). Bar = 500 μm. **(f, g)** Histograms showing total number of **(f)** corpora lutea, and **(g)** healthy growing follicles, counted across 100 serial sections of four ovaries collected from four animals at diestrus stage (mean ± SE; *P<0.05, t-test).

Histological analysis of ovaries, oviducts and uteri from 10-week-old virgin *Apc2^+/+^* and *Apc2^−/−^* mice revealed no gross morphological differences in the oviducts and uteri (representative images in Supplemental Figure 1). No problems were reported during labour in any of the experimental groups; it is therefore unlikely that uterine problems contribute to the observed subfertility phenotype. However, there was a significant decrease in the number of corpora lutea formed in *Apc2^−/−^* ovaries (Figure 1d, e & f), while the total number of growing follicles was increased, but not significantly (Figure 2d, e & g). Morphometric and histochemical analysis of corpora lutea did not reveal any histological differences in these structures between *Apc2^+/+^* and *Apc2^−/−^* ovaries (Supplemental Figure 2). Collectively, these findings suggest reduced ovulation is the cause of the subfertility observed in APC2-deficient mice.

**Figure 2:**
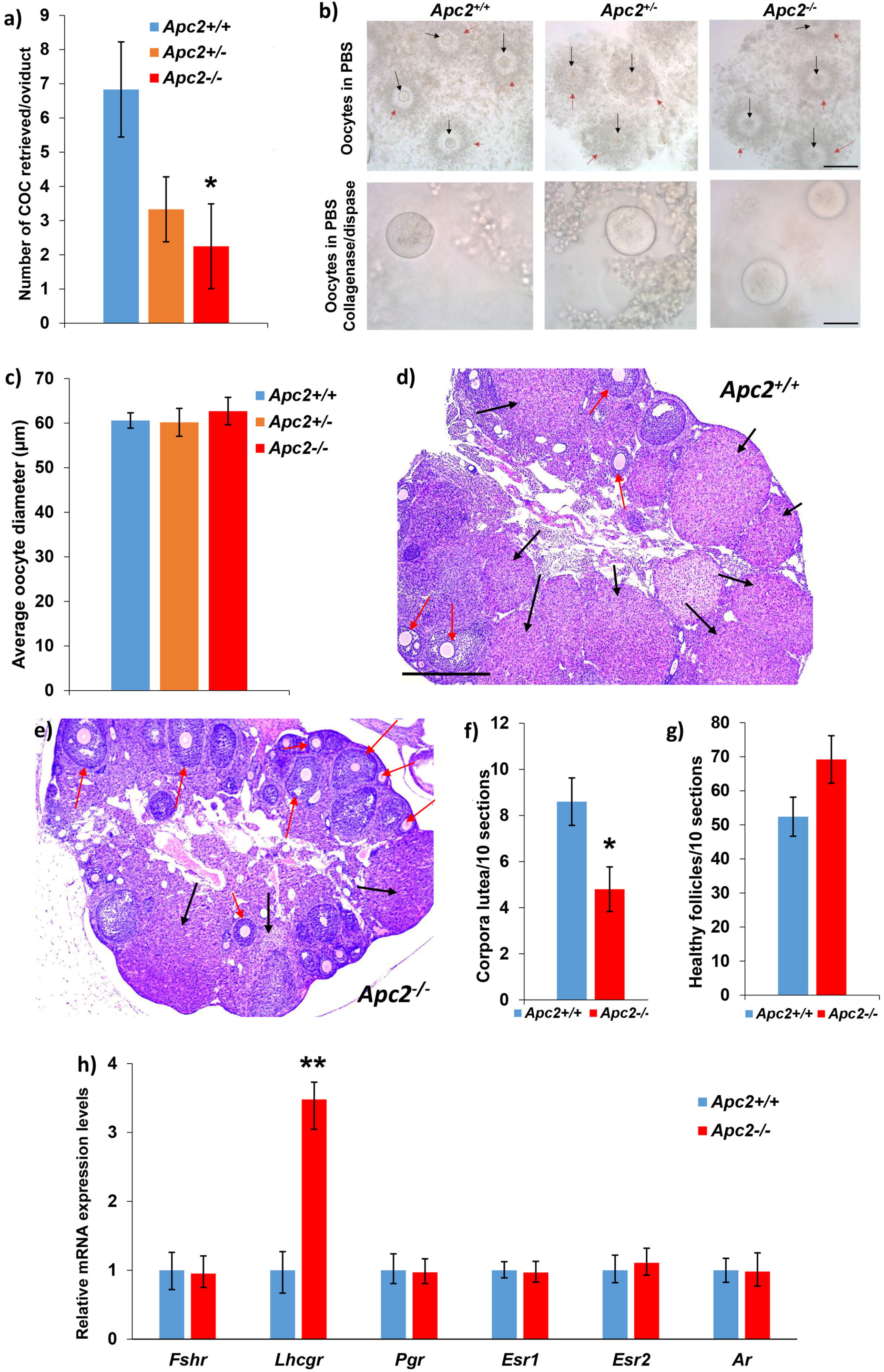
Exogenous gonadotrophin administration fails to reverse ovarian subfertility of APC2-deficient female mice. **(a)** APC2-deficiency caused a gene dose-dependent decrease in the number of ovulated COCs (mean±SE) retrieved from the oviducts post-superovulation. **(b)** Upper panels, representative photomicrographs of retrieved COCs showing the presence of oocytes (black arrows) surrounded by cumulus cells (red arrows). Bar = 200 μm. Lower panels, oocytes freed from cumulus cells. Bar = 50 μm. **(c)** Average oocyte diameter (mean±SE) among experimental groups, showing no difference. n=4 for *Apc2^+/+^*, n=3 for *Apc2^+/-^*, n=5 for *Apc2^−/−^*. Statistical significance between groups in panels a – c was determined using ANOVA test followed by LSD post hoc analysis (variances of experiment groups were homogeneous tested by Levene’s test). **(d, e)** Representative photomicrographs of **(d)** *Apc2^+/+^*, and **(e)** *Apc2^−/−^* superovulated ovaries, showing growing follicles (red arrows) and corpora lutea (black arrows). Bar = 500 μm. **(f, g)** Total number of **(f)** corpora lutea, and **(g)** healthy follicles counted across 100 serial sections of five superovulated stage-matched ovaries from different animals (mean ± SE; *P<0.05, t-test). **(h)** Gene expression levels of hormone receptors by qRT-PCR on RNA extracted from whole ovaries of *Apc2^+/+^* and *Apc2^−/−^* 10-week-old female mice. Relative expression levels are normalized to Actb expression. N=4 except for *lhcgr* and *Ar* in *Apc2^−/−^* where n=3 (mean ± 95% confidence intervals; **P<0.01, determined from confidence intervals) (70).

### Subfertility in APC2-deficient female mice is caused by intra-ovarian defects

Given the constitutive nature of the *Apc2* gene deletion in our mice, the genotype dose-dependent reduction in fertility, potentially as a result of an ovulation defect, may be due to defects in extra-ovarian regulation of ovarian function, triggered by hypothalamic/pituitary endocrine signals. To address this, follicle stimulating hormone (FSH) and luteinizing hormones (LH) levels in serum from 10-week old virgin *Apc2^+/+^* and *Apc2^−/−^* female mice at diestrus stage were analysed by ELISA, but showed no differences (Supplemental Figure 3a, b), suggesting hypothalamic/pituitary signals are not affected by *Apc2* deletion.

Next, to determine whether the response of the ovary to endocrine signals was compromised in the context of *Apc2* deletion, exogenous gonadotrophins were administered to induce superovulation in 10-week-old virgin *Apc2^+/+^, Apc2^+/-^* and *Apc2^−/−^* mice. There was a gene dose-dependent decrease in the number of cumulus oophorus complexes (COCs) collected from the ampulla post-superovulation (Figure 2a). However, morphological analysis of the COCs demonstrated that all oocytes were of comparable size, surrounded by a layer of cumulus cells of comparable thickness, and were healthy, with no signs of fragmentation, irrespective of genotype (Figure 2b, c).

Importantly, histological analysis post-superovulation demonstrated a significant reduction in the number of corpora lutea in super-ovulated *Apc2^−/−^* ovaries compared to *Apc2^+/+^* (Figure 2 d,e&f). As with unstimulated ovaries, a slight, but non-significant, increase in the number of healthy growing follicles in *Apc2^−/−^* ovaries was observed (Figure 2d,e&g). Taken together, these findings suggest that the subfertility phenotype seen in APC2-deficient female mice is not due to extra-ovarian defects in pituitary gonadotrophin secretion, but rather due to intra-ovarian defects in response to gonadotrophins that result in reduced ovulation. Therefore, expression levels of the ovarian gonadotrophin receptors *Fshr* and *Lhcgr*, together with the steroid hormone receptors *Pgr, Esr1, Esr2* and *Ar*, were assessed in ovaries from *Apc2^+/+^* and *Apc2^−/−^* mice. Significant over-expression of *Lhcgr* was evident in *Apc2^−/−^* ovaries (Figure 2h), but the other receptors were unaltered. Importantly, the LH receptor is a target of canonical WNT signalling (40), and its over-expression has previously been associated with infertility in mice (41).

Detailed morphometric analysis, on serial-sectioned ovaries collected at diestrus stage from *Apc2^+/+^* and *Apc2^−/−^mice*, demonstrated that the trend for an increase in the total number of healthy growing follicles in *Apc2:^−/−^* ovaries (Figure 1g), was restricted to the number of primary and antral follicles (Figure 3a). Size distribution analysis for healthy antral and preovulatory follicles demonstrated a significant increase in the percentage of small follicles (diameter <200μm) and a significant decrease in the percentage of larger follicles (diameter >300μm) in *Apc2^−/−^* ovaries (Figure 3b). Analysis of atretic follicles was undertaken to determine whether increased atresia was causing the reduction in larger follicles in *Apc2^−/−^* ovaries, but their total number and size distribution were not significantly altered (Supplemental Figure 3c,d). IHC for Ki67 revealed that proliferation was unaltered in follicular granulosa or theca cells (Supplemental Figure 3e). However, apoptosis, as measured by cleaved caspase 3 IHC, was significantly increased in granulosa cells in *Apc2^−/−^* follicles (Figure 3c,d&e). Histological analysis of pre-ovulatory follicles together with gene expression analysis of EFG ligands and receptor did not reveal defects in ovulation (Supplemental figure 3f,g). Thus, APC2 deficiency increases granulosa cell apoptosis, restricting follicular growth and reducing their ability to reach the pre-ovulatory stage.

**Figure 3:**
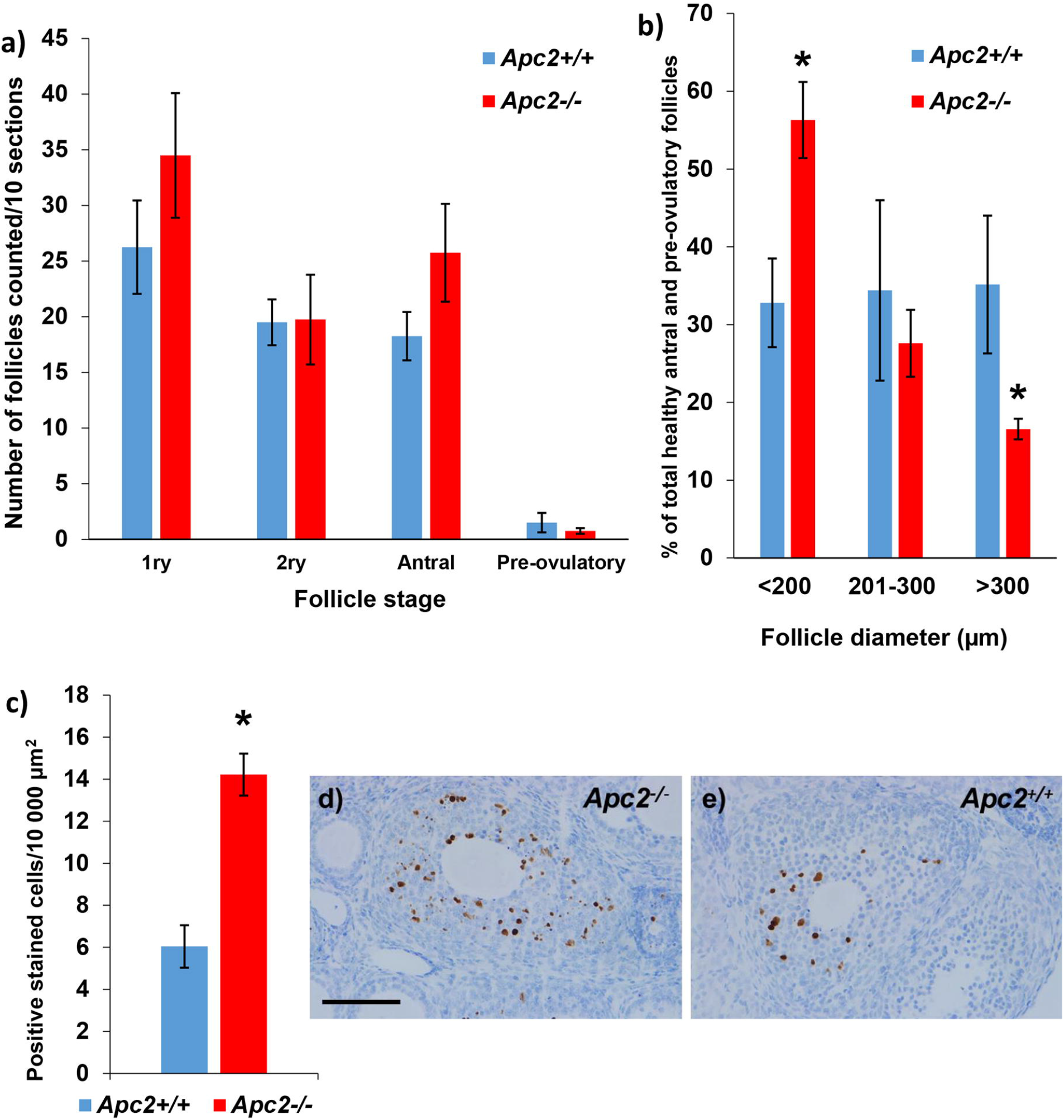
APC2-deficiency impairs follicular growth in the ovary. **(a)** Histogram showing total number of primary (1ry), secondary (2ry), antral and pre-ovulatory follicles in *Apc2^+/+^* and *Apc2^−/−^* ovarian sections (mean ± SE; n=4; no significant differences, t-test). **(b)** Size distribution of healthy antral and pre-ovulatory follicles (mean ± SE; n=4; *P<0.05, t-test). **(c)** Histogram showing a >2-fold increase of apoptosis in granulosa cells of *Apc2^−/−^* follicles (mean ± SE; n=4; *P<0.05, t-test). **(d, e)** Representative photomicrographs of cleaved caspase 3 immunostaining in (d) *Apc2^−/−^* and (e) *Apc2^+/+^* granulosa cells. Bars = 100 μm.

### APC2 deficiency activates ovarian WNT signalling and upregulates *Foxo1* expression

As APC2 is a known regulator of canonical WNT signalling, we investigated whether dysregulated WNT signalling was mechanistically linked to the restriction in follicular growth in *Apc2^−/−^* ovaries. Ovarian subcellular localization of β-catenin was assessed by immunohistochemistry (IHC), and expression of a standard panel of WNT target genes was determined by qRT-PCR, using whole ovaries collected at diestrus stage from 10-week-old virgin control (Apc2^+/+^) and APC2-knockout (*Apc2^−/−^*) mice. IHC analysis of β-catenin revealed a comparable pattern of expression in all ovarian compartments between control and knockout ovaries (Supplemental Figure 4), although increased staining intensity was notable in atretic follicles from *Apc2^−/−^* ovaries (Figure 4a). qRT-PCR analysis revealed a significant increase in the expression levels of *Apc, Axin2, Ctnnb1, Fgf1* and *Lgr5* in *Apc2^−/−^* ovaries compared to control ovaries (Figure 4b). However, there were no significant changes in *Cd44, Lef1* and *Wif1.*

**Figure 4:**
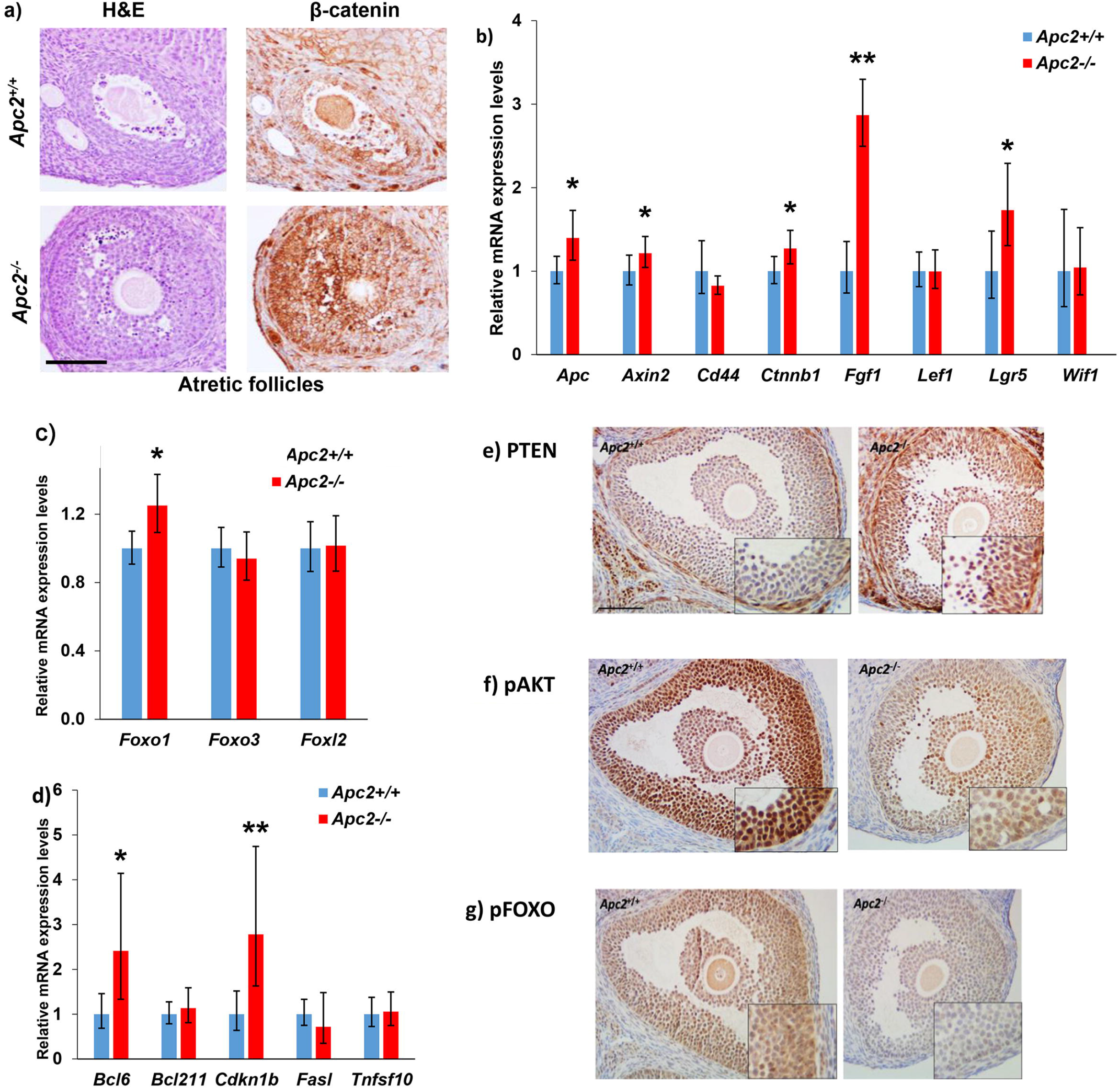
Identifying molecular mechanisms associated with subfertility in APC2-deficient ovaries. **(a)** Representative photomicrographs of H&E and β-catenin immunohistochemical staining of atretic follicles showing high expression of β-catenin in granulosa cells of atretic follicles in *Apc2^−/−^* vs. *Apc2^+/+^* ovaries. Bar = 100 μm. **(b)** Expression levels of a subset of WNT-target genes compared between *Apc2^+/+^* and *Apc2^−/−^* ovarian extracts by qRT-PCR. Expression levels of five out of eight genes analysed were significantly elevated in *Apc2^−/−^* ovaries (mean ± 95% confidence intervals; n =4; *P<0.05, **P<0.01, determined from confidence intervals) (70). **(c, d)** Relative expression levels of a panel of **(c)** *Fox* transcription factors, and **(d)** FOX downstream target genes in *Apc2^+/+^* and *Apc2^−/−^* 10-week-old ovaries (mean ± 95% confidence intervals; n=4 for all measurements except for *Fasl* and *Foxo1* in *Apc2^+/+^* where n=3; *P<0.05, **P<0.01, determined from confidence intervals) (70). **(e, f, g)** Representative photomicrographs of **(e)** PTEN, **(f)** p-AKT (ser-473), and **(g)** p-FOXO1,3,4 immunostaining in *Apc2^+/+^* and *Apc2^−/−^* ovarian follicles. Bars = 200 μm.

Given the established role of the FOX family of transcription factors as regulators of apoptosis within ovarian granulosa cells (42–44), and their increased expression in granulosa cells of cultured follicles post-WNT signalling activation (45, 46), gene expression levels for *Foxo1, Foxo3* and *Foxl2* were analysed in *Apc2^+/+^* and *Apc2^−/−^* whole ovaries. A significant increase in *Foxo1* expression levels was seen in *Apc2^−/−^* ovaries (Figure 4c). Furthermore, the FOXO target genes *Bcl6* and *Cdkn1b* were significantly upregulated in *Apc2^−/−^* ovaries compared to controls (Figure 4d).

The PTEN/PI3K/AKT signalling pathway is an established regulator of FOXO transcriptional activity and post-translational modification (47). On activation of AKT, FOXO proteins are inactivated by phosphorylation and translocated from nucleus to cytoplasm (47). In addition, the crosstalk between activated WNT signalling and PTEN, causing the over-expression of the latter, is well established (16, 17, 48). IHC analysis of PTEN, p-AKT and p-FOXO 1,3,4 in *Apc2*^+/+^ and *Apc2*^−/−^ ovaries revealed that PTEN expression was stronger in theca and granulosa cells of *Apc2^−/−^* follicles (Figure 4e). This was accompanied by a reduction in p-AKT immunostaining in *Apc2^−/−^* granulosa cells (Figure 4f) and a consequent reduction in p-FOXO1,3,4 levels (Figure 4g). Thus, the increased apoptosis seen in *Apc2^−/−^* follicles is likely due to upregulation of *Foxo1* and its downstream effector genes, secondary to decreased activation of PI3K/p-AKT signalling caused by PTEN upregulation.

### APC2-deficient ovaries show impaired vascularisation and steroidogenesis

Interaction between β-catenin and FOXO1 has been previously described to affect tight junctions in endothelial cells disrupting angiogenesis (49). Follicular growth impairment has been shown to occur following angiogenesis disruption, because the vascular network surrounding the growing follicles is essential for follicular development (50). IHC for CD34 was used to compare follicular vascularization between *Apc2^+/+^* and *Apc2^−/−^* ovaries. While late antral/pre-ovulatory follicles in *Apc2^+/+^* ovaries were surrounded by 2 continuous layers of endothelial cells, those in *Apc2^−/−^* mice showed discontinuous layers (Figure 5a). Furthermore, a significantly reduced level of *Vegfa* expression (Figure 5b) supports the notion of impaired vascularisation within *Apc2^−/−^* ovaries, although it could, in part, be attributed to the reduced number of corpora lutea.

**Figure 5:**
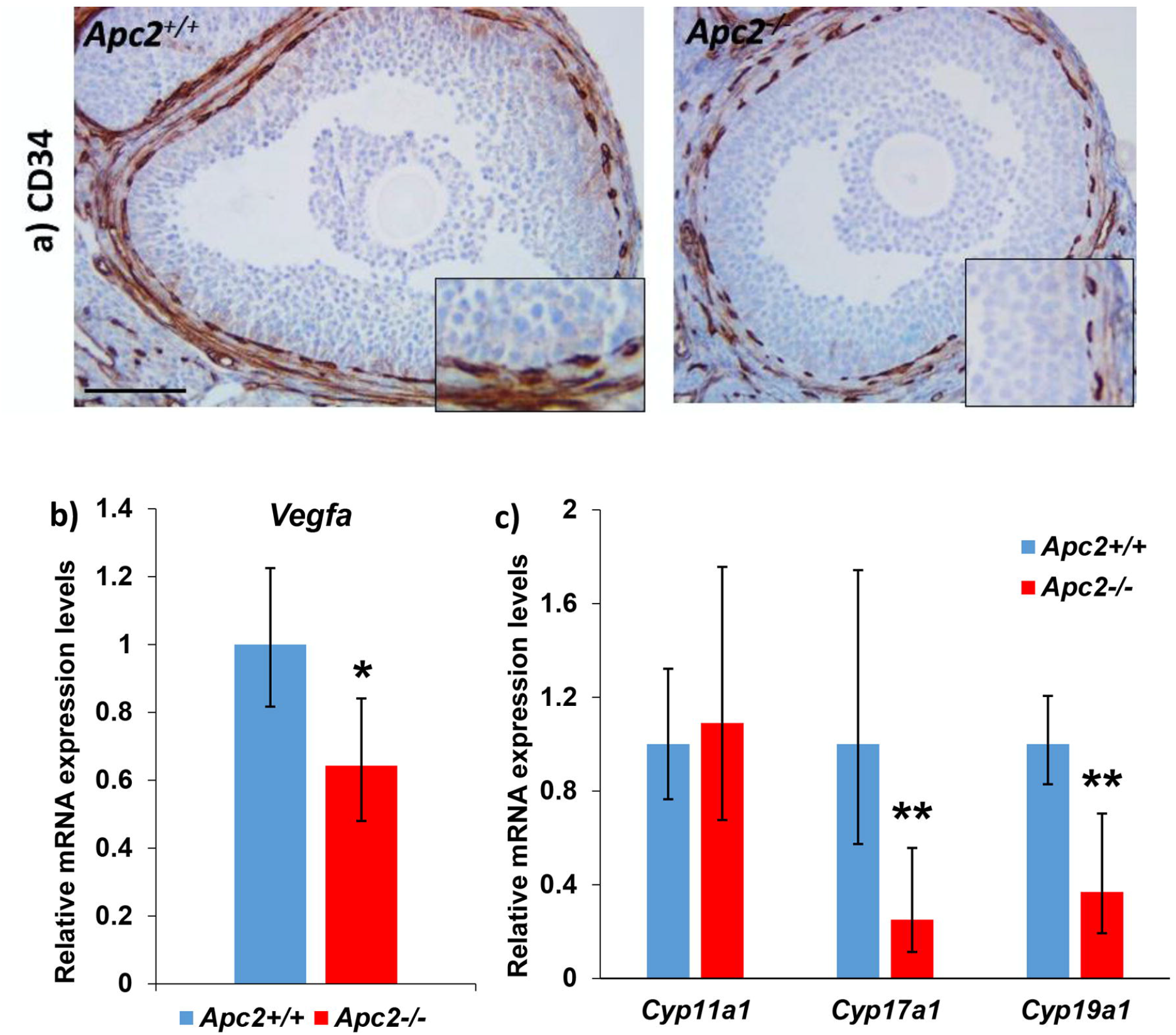
APC2 loss impairs follicle steroidogenesis and vascularization. **(a)** Representative photomicrographs of CD34 immunostaining of follicles in *Apc2*’\** and *Apc2^−/−^* 10-week-old ovaries. Bars = 100 μm. Insets are magnified 2X. **(b, c)** Gene expression levels of **(b)** *Vegfa*, and **(c)** steroidogenic enzymes by qRT-PCR on ovaries of 10-week-old Apc2+/+ and Apc2-/-female mice (mean ± 95% confidence intervals; n=4 for all measurements except for *Cyp19a1* in *Apc2^−/−^* where n=3; *P<0.05, **P<0.01, determined from confidence intervals) (70).

Negative regulation of follicle steroidogenesis by canonical WNT signalling has also previously been demonstrated (46). We therefore examined the expression of key enzymes required for steroidogenesis in *Apc2* knockout ovaries. We found there was significantly reduced expression of both *Cyp17a1* (coding for steroid 17-α-hydroxylase/17,20 lyase) and *Cyp19a1* (coding for aromatase) in *Apc2^−/−^* ovaries compared to *Apc2^+/+^* ovaries (Figure 5c).

Therefore, activation of WNT signalling in *Apc2* knockout ovaries results in overexpression of PTEN and a reduction in activity of steroidogenesis and angiogenic pathways. These metabolic defects combine to result in a reduced number of follicles maturing to the ovulatory stage.

### Long-term activation of WNT signalling by APC/APC2 deficiency, results in ovarian adult granulosa cell tumour formation

Because of the WNT signalling-dependent defects observed in 10-week-old *Apc2* knockout mice, and the potential role of WNT signalling in driving ovarian tumour development in mice (15, 17, 48, 51, 52), we aged cohorts of mice in which WNT signalling was activated to different levels using a hypomorphic *Apc^fl/fl^* (weak), a hypomorphic *Apc^fl/fl^* plus *Apc2^+/−^* knockout (moderate) or a hypomorphic *Apc^fl/fl^* plus *Apc2^−/−^* knockout (strong). No gross ovarian tumours were detected in any cohorts at 6 months of age; however 6/29 (20.7%) of the APC2-deficient cohorts *(Apc2^+/−^* and *Apc2^+/−^* cohorts on the background of *Apc^fl/fl^)* had developed adult ovarian GCTs at 12-18 months of age as compared to 1/19 (5.26%) of the APC2-proficient *(Apc^fl/fl^)* cohort developing ovarian GCT at 18 months of age (Table 1).

**Table 1:** Frequency of GCT formation in 12 and 18-month-old *Apc2* experimental genotypes on the background of *Ap^fl/fl^*. *One animal developed bilateral tumours.

| Age (months) | Genotype | Frequency of ovarian GCT |
| --- | --- | --- |
| 12 | *Apc2*<sup>+/+</sup> | 0/10 (0%) |
| 12 | *Apc2*<sup>+/-</sup> | 0/4 (0%) |
| 12 | *Apc2*<sup>-/-</sup> | 2*/10 (20%) |
| 18 | *Apc2*<sup>+/+</sup> | 1/9 (11.1%) |
| 18 | *Apc2*<sup>+/-</sup> | 3/9 (33.3%) |
| 18 | *Apc2*<sup>-/-</sup> | 1/6 (16.67%) |

The tumours ranged from small microscopic *in situ* tumours to large macroscopic tumours (Figure 6a-e). Morphologically, they recapitulated human ovarian adult GCTs and showed a range of different histological patterns (Figure 6f-l). Cells were highly anaplastic (Figure 6m) and mitotic figures were evident (Figure 6n). Call-Exner bodies (formed of follicle remnants, Figure 6j,o) and coffee bean-shaped nuclei (Figure 6o), both characteristic of adult GCTs, were occasionally present.

**Figure 6:**
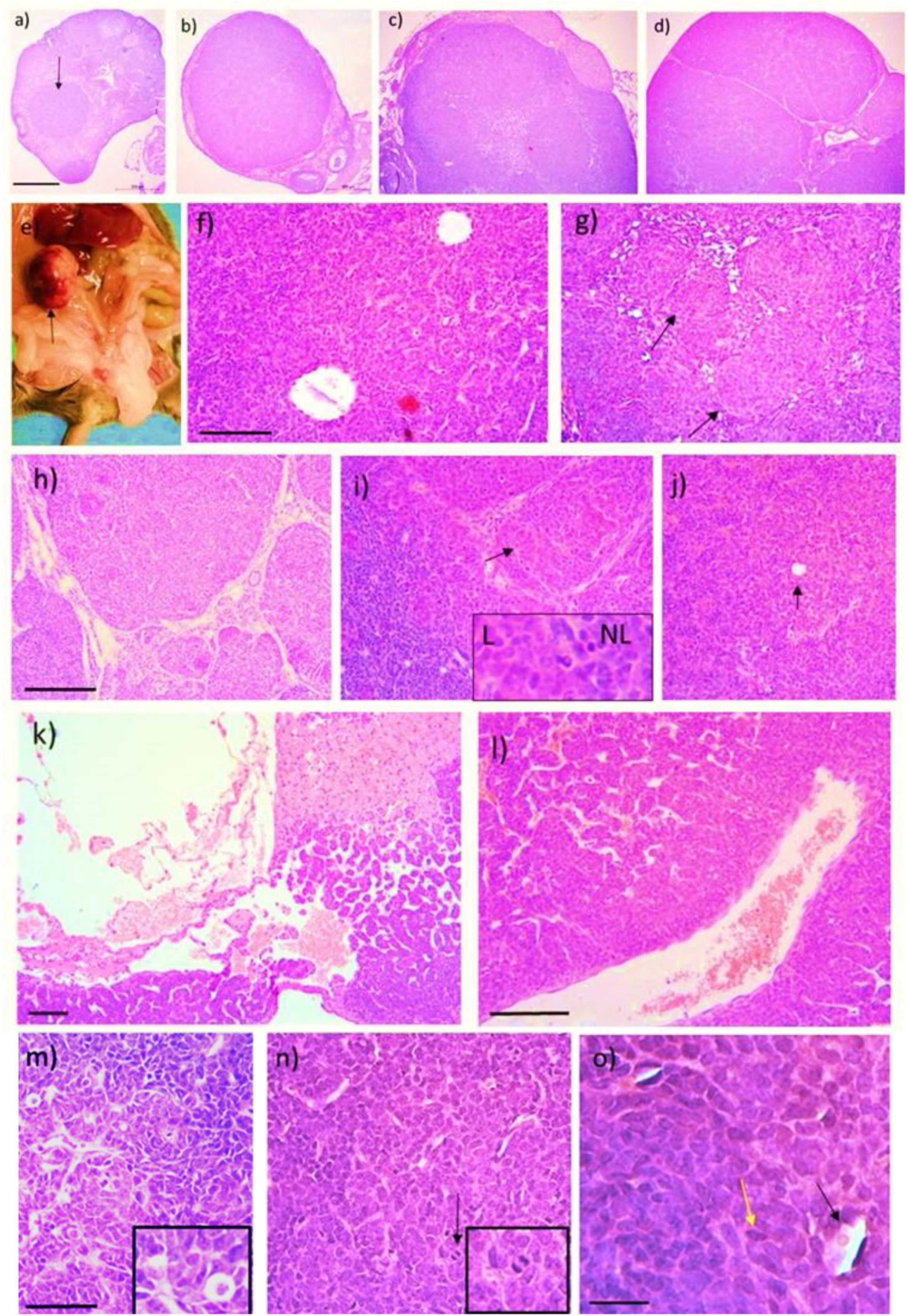
Aging APC2-deficient mice develop adult GCTs. Tumours ranged in size from **(a)** small *in situ* tumours (arrow) to **(b)** small tumours of normal ovarian size, **(c, d)** small but macroscopically visible tumours or **(e)** a large macroscopic tumour (arrow). The tumours displayed varying histologic patterns such as **(f)** follicular, **(g)** nodular (arrows), **(h)** insular, **(i)** luteinized (arrow: luteinized area shown at 4X original magnification in inset, L: luteinized, NL: non-luteinized), **(j)** diffuse (arrow: Call-Exner body), and **(k)** cystic patterns. The tumours were **(l)** highly vascularized, **(m)** anaplastic, **(n)** mitotic (black arrow), and showed **(o)** Call-Exner bodies (black arrow) and coffee-bean nuclei (yellow arrow). Bars a-d = 500 μm, f-l = 100 μm, m-n = 50 μm, o = 20 μm.

The molecular signature of the tumours was assessed by IHC for markers associated with human adult GCTs. Several studies have shown increased FOXL2 protein expression in human GCTs and animal models (53–55), and in agreement with these findings, our APC2-deficient GCTs also displayed elevated levels of FOXL2 IHC expression (Figure 7a, Supplemental Figure 5a). Human ovarian GCTs are also characterized by frequent focal staining for estrogen receptor alpha (ERα), and the staining seen in our tumours followed this pattern (Figure 7b, Supplemental Figure 5b). Immunohistochemistry for inhibin has been previously used in the differential diagnosis of GCTs (56). APC2-deficient GCTs were positive for inhibin-α IHC, which showed focal cytoplasmic staining (Figure 7c).

**Figure 7:**
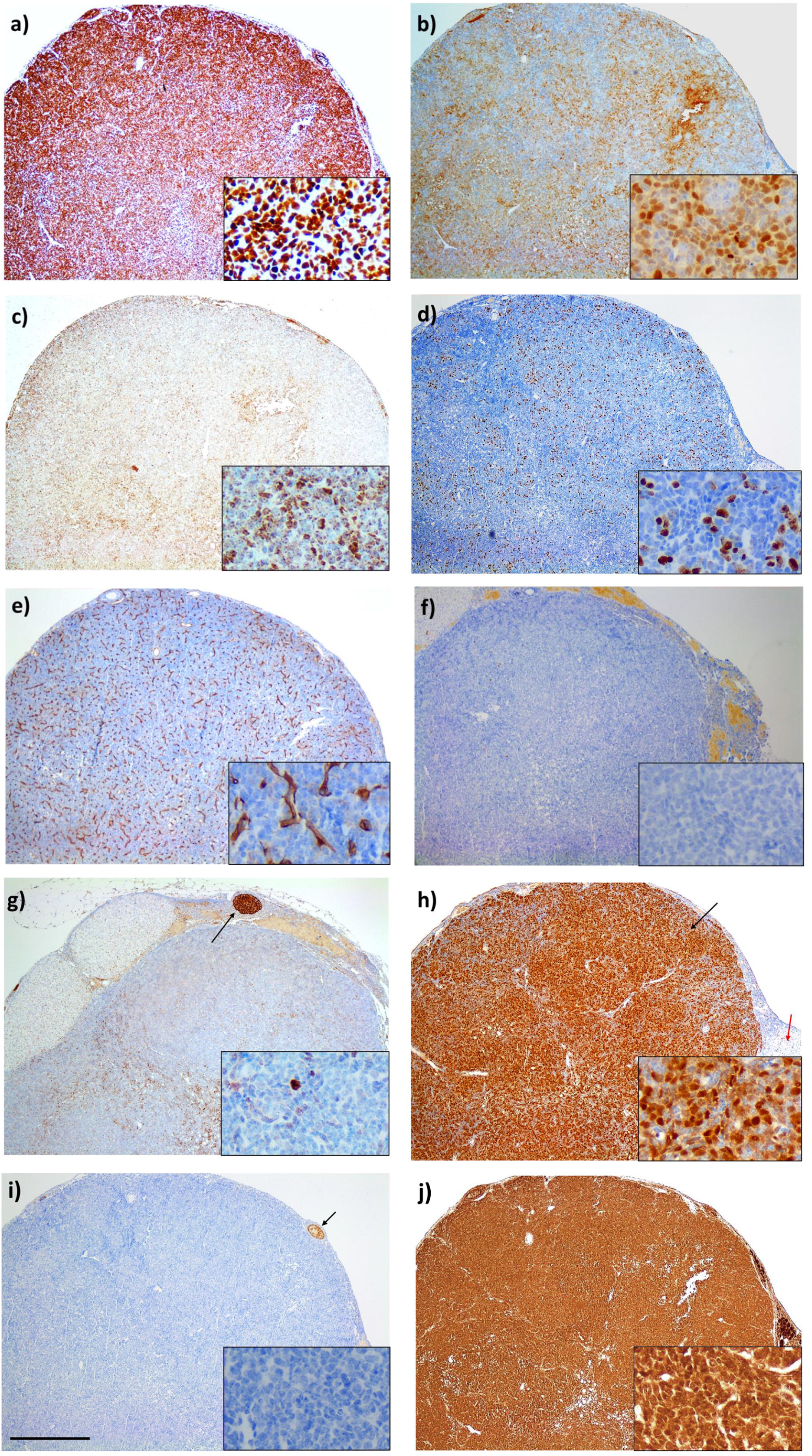
Molecular characterization of GCTs developing in 18-month *Apc2^+/−^* ovaries. Representative photomicrographs of immunohistochemical staining for **(a)** FOXL2 protein **(b)** estrogen receptor alpha (ER), **(c)** Inhibin-, **(d)** Ki67 **(e)** CD34 **(f)** cleaved caspase-3 (showing absence of apoptosis in GCT), (g) FOXO1 showing weak staining in GCT, in contrast to a growing follicle (black arrow), (h) β-catenin showing intense staining in GCT (black arrow) as compared to a corpus luteum (red arrow), **(i)** p-AKT which was absent in GCT as compared to a growing follicle (black arrow), and **(j)** PTEN. Bar = 500 μm. Insets are 5X original magnification.

IHC for Ki67, CD34 and cleaved caspase 3 demonstrated the classic hallmarks of proliferation, neovascularization (angiogenesis) and absence of apoptosis respectively, in all GCTs analysed (Figure 7d-f). As previously mentioned, impaired follicular growth in *Apc2^−/−^* ovaries was associated with increased apoptosis (Figure 3c-e) and *Foxo1* expression (Figure 4c). As apoptosis was reduced in APC2-deficient GCTs, we analysed FOXO1 expression levels by IHC and saw a reduction in FOXO1 staining in GCT area as compared to granulosa cells of growing follicles (Figure 7g, Supplemental Figure 5c).

To determine whether active canonical WNT signalling was associated with GCT formation, β-catenin IHC was undertaken on one tumour collected from the 12-month-old *Apc2:^1^*cohort, one tumour from the 18-month-old *Apc2^+/+^* cohort and 3 tumours form the 18-month-old *Apc2^+/-^* cohort. In all cases, the tumour areas (Figure 7h, black arrows) strongly expressed γ-catenin in contrast to the expression seen in non-tumour areas (Figure 7h).

Due to limitation of available tumour samples, qRT-PCR analysis was performed on a subset of WNT signalling target genes using RNA from two Apc2-deficient GCTs, with *Apc2-* proficient ovaries used as control material. Two independent areas from each tumour were analysed to allow for tumour heterogeneity. Comparison of expression levels demonstrated higher levels of *Wif1* and *Axin2* within APC2-deficient tumours (Supplemental Figure 5d,e).

Activation of PI3K/AKT signalling via *Pten* deletion has been shown to enhance GCT development and progression in alternative mouse models driven by Wnt signalling activation (57, 58). However, IHC for phospho-AKT (p-AKT), a marker of active PI3K/AKT signalling, revealed undetectable levels within our APC2-deficient tumours (Figure 7i). The lack of p-AKT is most likely due to the highly-elevated levels of PTEN within the tumours (Figure 7j).

Thus, APC2 deficiency, when combined with a hyopomorphic *Apc^fl^* allele, resulted in a higher propensity for the formation of ovarian GCTs with histological and molecular features characteristic of human adult GCTs.

## Discussion

This study has revealed, for the first time, that APC2-deficiency activates WNT signalling in the ovary during early adulthood, which subsequently disrupts ovarian homeostasis and causes subfertility originating from an ovarian defect. Follicle growth was perturbed in APC2-deficient mice secondary to defective response to gonadotrophins, reduced follicular vascularity, downregulation of genes coding for steroidogenic enzymes and upregulation of *Foxo1* expression, which contributed to increased apoptosis of granulosa cells in APC2-deficient follicles. We have also shown at least 20% of APC2-deficient female mice (on the background of non-induced *Apc^fl/fl^*) develop WNT-driven GCT as early as 12 months. These tumours recapitulated human adult GCT histology and molecular features.

Our findings highlight the role of APC2 as an important regulator of WNT signalling in the ovary. Although initial studies performed in *Drosophila* and on cell lines to functionally-characterize APC2 demonstrated the presence of β-catenin and Axin1 binding sites in APC2, which enable it to regulate WNT signalling (23, 27, 59–62), in an *in vivo* mammalian setting, APC2 function is tissue specific. APC2 loss in the mouse small intestine and liver resulted in activation of WNT signalling but not in the mammary glands (34, 35). Hence, the functions of APC2 cannot be extrapolated from one mammalian system to another without direct experimentation.

The current findings also extend our knowledge of deleterious effects of WNT signalling activation on ovarian homeostasis and fertility (10, 15–17). We have shown that reduced ovulation observed in APC2-deficient mice is not caused by defects in ovulation and terminal differentiation of granulosa cells (which happen when WNT signalling is activated in antral follicles), but rather caused by restricted follicular growth and failure to reach the preovulatory stage. This phenotype is similar to previous phenotypes published when WNT signalling was activated in pre-antral follicles (16, 17), implying that APC2 activity is required in growing follicles as early as the pre-antral stage.

The tumour suppressor role of APC2 protein in ovarian granulosa cell tumour formation has also been highlighted for the first time and the current study provides further evidence of the roles of WNT signalling activation in the pathogenesis of ovarian GCT. These findings build on previous work pointing to this role of WNT signalling in clinical data (17, 19, 20, 63), and in GEMMs (15, 17) but as noted above, given the tissue-specific effects of APC2 knockout, could not have been predicted *a priori.*

Given the constitutive nature of the *Apc2* null allele, both autonomous and non-autonomous mechanisms are expected to contribute to the phenotypes described. Results of the current study have clearly shown the intra-ovarian origin of the subfertility phenotype described in APC2-deficient mice, and that hypothalamic-pituitary regulation of ovarian function is not contributing to the subfertility phenotype. Although the subfertility is caused by increased apoptosis of granulosa cells, a contribution of endothelial cells to the phenotype was evident. Whether the same phenotype could be reproduced if APC2-deletion was targeted exclusively to granulosa cells (e.g. using *Amhr2* or *Cyp19a-cre)* remains unknown, due to the unavailability of an *Apc2* conditional allele. The same applies to GCTs developing in APC2-deficient mice, which – in contrast – displayed enhanced angiogenesis.

Also, it is unlikely that WNT signalling activation is the sole driver of the reported phenotypes and cross talk between WNT signalling and other signalling pathways must also be considered. For example, unlike in early adulthood, FOXO1 expression was absent in APC2-deficient GCT, implying a need to silence FOXO1 and to stop FOXO1-driven granulosa cell apoptosis as a prerequisite for tumourigenesis. It has been previously shown that knocking out *Foxo1/Foxo3* leads to the development of GCT in 20% of female mice (54). However, the cause of the ‘switch’ from FOXO1 being present and granulosa cell apoptosis to absent FOXO1 with granulosa cell proliferation and tumourigenesis was not identified and needs to be further characterized. The high levels of PTEN in granulosa cells of growing follicles might have contributed to increased apoptosis by inhibiting the translocation of FOXO1 outside the nucleus and thus ensuring FOXO1 activates pro-apoptotic target genes. In addition, high PTEN expression levels found in GCT of APC2-deficient ovaries might be responsible for the late development of tumourigenesis, as previously described in other models (Lague *et al.* 2008; Richards *et al.* 2012; Liu *et al.* 2015). It is thus possible to hypothesize that, similar to previously published models, deleting *Pten* in granulosa cells of APC2-deficient ovaries would lead to rapid tumour development.

This study has caveats. One limitation was that the breeding data available for different genotypes of female *Apc2* mice *(Apc2^+/+^, Apc2^+/−^*, *Apc2^−/−^*) represented crossings to males of the corresponding genotype, rather than to wild type males. Effects of Apc2-gene dosage on male fertility are not yet characterized, with the caveat that male fertility might be affected in APC2-deficient male mice, and could contribute to the delayed pregnancy and reduced litter size observed in APC2-deficient crosses. However, retrieval and counting of ovulated oocytes post-gonadotrophin administration confirmed that APC2-deficient female mice ovulate less and would be expected to give smaller litter size. Impairment of response to gonadotrophin is mediated by overexpression of *Lhcgr*, which has been recently reported to cause complete infertility in female mice, with histological analysis revealing that follicles failed to progress beyond the pre-antral stage (41). Over-expression of *Lhcgr* in APC2-deficient mice most likely occurs due to canonical WNT signalling activation, as a 3.5-fold increase in *Lhcgr* expression levels has been reported in granulosa cells transduced with constitutively-active β-catenin, in the presence of FSH (40). In addition, this early elevation of *Lhcgr* expression might have contributed to GCT development (41). Another important caveat to this study was the small numbers of aged *Apc2^−/−^* mice available for tumour development studies. This was, unfortunately, an unavoidable consequence of the reduced fertility phenotype in these animals.

In conclusion, this study advances our understanding of the role of WNT signalling in ovarian homeostasis and tumourigenesis, and of the role played by APC2 in regulating this pathway. The finding that WNT signalling activation in growing follicles impairs ovulation raises the importance of the assessment of WNT signalling activation in the setting of human female subfertility/infertility. This could provide new insights into the molecular pathogenesis of this condition, and may help in designing new treatment interventions for these patients. Furthermore, our findings extend the list of mutations which cause female subfertility or infertility in early adulthood in mice followed by development of GCT upon aging (15, 17, 41, 54). It remains to be determined if a similar link exists in humans and, if so, what are the molecular drivers, but APC2 must now be included on the list of candidates which should be investigated in this clinical context.

## Materials and Methods

### Animal models, fertility and ovulation rate

All experiments were carried out under the authority of UK Home Office personal and project licences and according to ARRIVE guidelines. Genotyping for the constitutive knockout allele of *Apc2 (Apc2^-^)* and the hypomorphic allele of *Apc (Apd^fl^)* (35, 37–39) were performed as previously described (35, 37) (Supplementary Table 1).

To assess female fertility, retrospective analysis of breeding performance was analysed from cages in which two 7-11 week-old female mice of the experimental genotypes *(Apc2^+/+^, Apc2^+/−^* and *Apc2^−/−^*) were housed with a 7-9 week-old male of the same genotype for 3 months (n=4 cages). Litter sizes were determined at the time of weaning.

To determine ovulation rates, 10 week-old female mice were super-ovulated by a single intraperitoneal injection of 5 IU pregnant mare’s serum gonadotropin (MSD animal health, UK), followed by 5 IU human chorionic gonadotrophin (MSD animal health, UK), 47 hours later (64). Mice were either culled 16-17 (for COC retrieval) (65) or 22-24 hours later (for histological analysis).

### COC retrieval and characterization

After release from the oviducts, COCs were counted and examined by bright-field microscopy to assess morphology. Oocytes were freed from surrounding cumulus cells by addition of 40μl of 4mg/ml collagenase/dispase (Roche, Switzerland), dissolved in DMEM/F12 medium (Mediatech, USA), for 10 minutes, and examined to determine their integrity (66) and to measure their diameter (67).

### Histological analysis of ovaries

Follicle counting was performed on ovaries from 10-week-old *Apc2^+/+^* and *Apc2^−/−^* mice, either from randomly cycling females staged manually (using the vaginal cytology method) and collected at diestrus stage (n=4) or 22-24 hours post HCG administration (n=5). Each ovary was serially-sectioned into 100 5μM sections and each 10^th^ section was stained with H&E. Growing follicles were counted every 10^th^ section, when an oocyte nucleus was visible. Identification and classification of growing follicles and atretic follicles were performed as previously described (68, 69). The total number of follicles throughout the 10 counted sections was used. Follicle sizes were measured using a minimum of 4 diameters/follicle.

### Hormonal analysis

Serum hormonal levels were measured in 10-week-old *Apc2^+/+^* and *Apc2^−/−^* mice at diestrus stage using ELISA kits for FSH (Novateinbio, USA) and LH (Enzo Lifesciences, UK).

### Immunohistochemistry

Tissue sectioning and immunohistochemistry were performed as previously described (35), using primary antibodies listed in Supplementary Table 2. Sections were examined with an Olympus BX43 light microscope and microphotographs taken using a 5 Megapixel HD Microscope Camera (Leica MC170 HD, Germany).

### Quantitative RT-PCR analysis

RNA was extracted from whole ovaries or tumour pieces using RNeasy Plus mini extraction kit (Qiagen, Germany) and reverse transcription performed using QuantiTect Reverse transcription kit (Qiagen, Germany). All quantitative real time rtPCR assays were carried out three times using TaqMan^®^ universal master mix II with UNG (Applied Biosystems, USA), Taqman^®^ assays (Supplementary table 3) and QuantStudio™ 7 Flex Real Time PCR system (ThermoFisher, USA), and relative expression levels determined using QuantStudio™ 7 Real Time PCR software.

### Statistical analysis

Statistical significance for qRT-PCR data was determined from 95% confidence intervals (70). All other statistical analyses were performed using IBM SPSS version 20 (SPSS Inc, Chicago, IL, USA). Significance testing was performed using 2-tailed Student’s t-test, when 2 experimental cohorts were compared. When more were compared, ANOVA test was used, followed by a post-hoc test (LSD or Games-Howell). P-values of <0.05 were considered statistically significant.

## Acknowledgements

This project was funded by Egyptian Ministry of Higher Education (represented by the Egyptian Educational Bureau in London) and Cancer Research UK (ARC Programme grant C1295/A15937). We thank Professor Hans Clevers for providing us with *Apc2^−/−^* mouse, Professor Owen Sansom for critically reviewing the results, Professor Geraint Williams for training and guidance on histopathological assessment of ovaries, Elaine Taylor for assistance with mouse husbandry, Mark Bishop and Matthew Zverev for technical support and genotyping and Derek Scarborough for histologic preparation of tissues.

## Author Contributions

The research project was designed by NM, TH, MJS and ARC. NM, TH and KRR managed the mouse intercrosses. NM performed all data collection and analysis. The manuscript was drafted by NM and KRR. This manuscript is dedicated to the memory of the late Professor Alan Clarke. All other authors critically reviewed the manuscript and approved the final version submitted.

## Summary of Supplementary Information

Seven files:

Supplementary Tables 1 – 3 (.doc):

Supplementary Table 1: Primer sequences and reaction conditions used in genotyping.

Supplementary Table 2: Primary antibodies for immunohistochemistry.

Supplementary Table 3: Taqman assays used for relative gene expression analysis.

Supplementary Figure Legends (.doc)

Supplementary Figures:

Supplementary Figure 1: APC2 is dispensable for oviduct and uterine gross morphology (.jpg).

Supplementary Figure 2: APC2 is dispensable for corpora lutea (.jpg).

Supplementary Figure 3: Constitutive loss of APC2 has no effect on fertility hormones produced by pituitary gland or on the ovulation process (.jpg).

Supplementary Figure 4: Immunohistochemical localization of β-catenin protein in ovaries of *Apc2^+/+^* and *Apc2^−/−^* 10-week-old female mice (.jpg).

Supplementary Figure 5: Gene expression analysis of GCTs formed in a subset of APC2-deficient ovaries (.jpg).

