## Supplementary Figure Legends for "APC2 is Critical for Ovarian WNT Signalling Control, Fertility and Tumour Suppression"

**Supplementary Information**

**Supplementary Figure Legends**

**Supplementary Figure 1: APC2 is dispensable for oviduct and uterine gross morphology.** Microphotographs of **(a,b)** oviducts and **(c,d)** uteri from **(a,c)** *Apc2^+/+^* and **(b,d)** *Apc2^-/-^* 10-week-old female mice collected at diestrus stage. Bars = 100 µm for oviducts and 200 µm for uteri.

**Supplementary Figure 2: APC2 is dispensable for corpora lutea. (a,b)** Representative photomicrographs of corpus luteum in **(a)** *Apc2*^+/+^, and **(b)** *Apc2*^-/-^ ovarian sections. **(c,d)** corpora lutea size **(c)**, and Ki67 cell counts **(d)** in the two genotypes (mean ± SE; n = 4; no significant differences, t-test). **(e,f)** Representative photomicrographs of Ki67-positive cells (black arrows) in **(e)** *Apc2*^+/+^ and **(f)** *Apc2*^-/-^ corpora lutea. **(g,h)** Absence of cleaved caspase 3 staining in corpora lutea of **(g)** *Apc2*^+/+^, and **(h)** *Apc2*^-/-^ ovarian sections. Positive cleaved caspase 3 staining, as a control, is indicated by red arrows in atretic follicles. **(i, j)** Representative photomicrographs of CD34 staining in corpora lutea of **(i)** *Apc2*^+/+^ and **(j)** *Apc2*^-/-^ ovaries. Bars = 200 µm.

**Supplementary Figure 3: Constitutive loss of APC2 has no effect on fertility hormones produced by pituitary gland or on the ovulation process. (a,b)** Serum levels of **(a)** FSH and **(b)** LH measured by ELISA (mean ± SE; n = 4 for LH, 6 for FSH; no significant differences, t-test). **(c)** Quantitation of atretic follicles in ovaries of *Apc2*^+/+^ and *Apc2*^-/-^ mice (mean ± SE; n = 4 no significant differences, t-test). **(d)** Size distribution of atretic antral and pre-ovulatory follicles (mean ± SE; n = 4; no significant differences, t-test). **(e)** Comparable proliferation of *Apc2*^+/+^ and *Apc2*^-/-^ follicular cells (mean ± SE; n = 4; no significant differences, t-test). **(f)** Representative photomicrograph of follicles approaching the pre-ovulatory stage in *Apc2*^+/+^ and *Apc2*^-/-^ ovarian sections. Bar = 100 µm. **(g)** qRT-PCR gene expression levels of a subset of *Egf* ligands and the *Egfr* receptor, known to regulate COC expansion, in *Apc2*^+/+^ and *Apc2*^-/-^ ovarian extracts (mean ± 95% confidence intervals; n=4; **P<0.01, significance determined from confidence intervals) (Cumming *et al.* 2007).

**Supplementary Figure 4: Immunohistochemical localization of β-catenin in ovaries.** **(a-g)** 10-week-old female *Apc2*^+/+^ mice. **(h-n)** 10-week-old female *Apc2*^-/-^ mice. OSE: ovarian surface epithelium. 1ry: primary follicle. 2ry: secondary follicle. SA: small (early) antral follicle. LA: large antral follicle. PO: pre-ovulatory follicle. Cl: corpus luteum. AF: atretic follicles. Bars = 100 µm, except **(f, l)** bars = 200 µm. Insets shows granulosa cells of growing follicles at 4X original magnification.

**Supplementary Figure 5: Gene expression analysis of GCTs formed in a subset of APC2-deficient ovaries.** qRT-PCR analysis of gene expression levels in an APC2 null tumour from a 12-month old animal and one APC2 heterozygote tumours from 18-month old animals compared to age-matched APC2 wild type ovaries (n=3 at both ages). Expression levels of GCT molecular markers *Foxl2* **(a)** and *Esr1* **(b)** were elevated in tumours while *Foxo1* expression **(c)** was reduced. Negative regulators of WNT signalling *Wif1* **(d)** and *Axin2* **(e)** were strongly increased in tumours. n=3 for 12-months and 18-months APC2 wild type ovaries. Two independent samples from the each tumour were analysed and results from both these samples are shown. The small numbers of tumour samples available means that statistical analysis is not possible. T: Tumour.
