## Supplementary figures and images for "APC2 is Critical for Ovarian WNT Signalling Control, Fertility and Tumour Suppression"

### Supplementary Figure 1

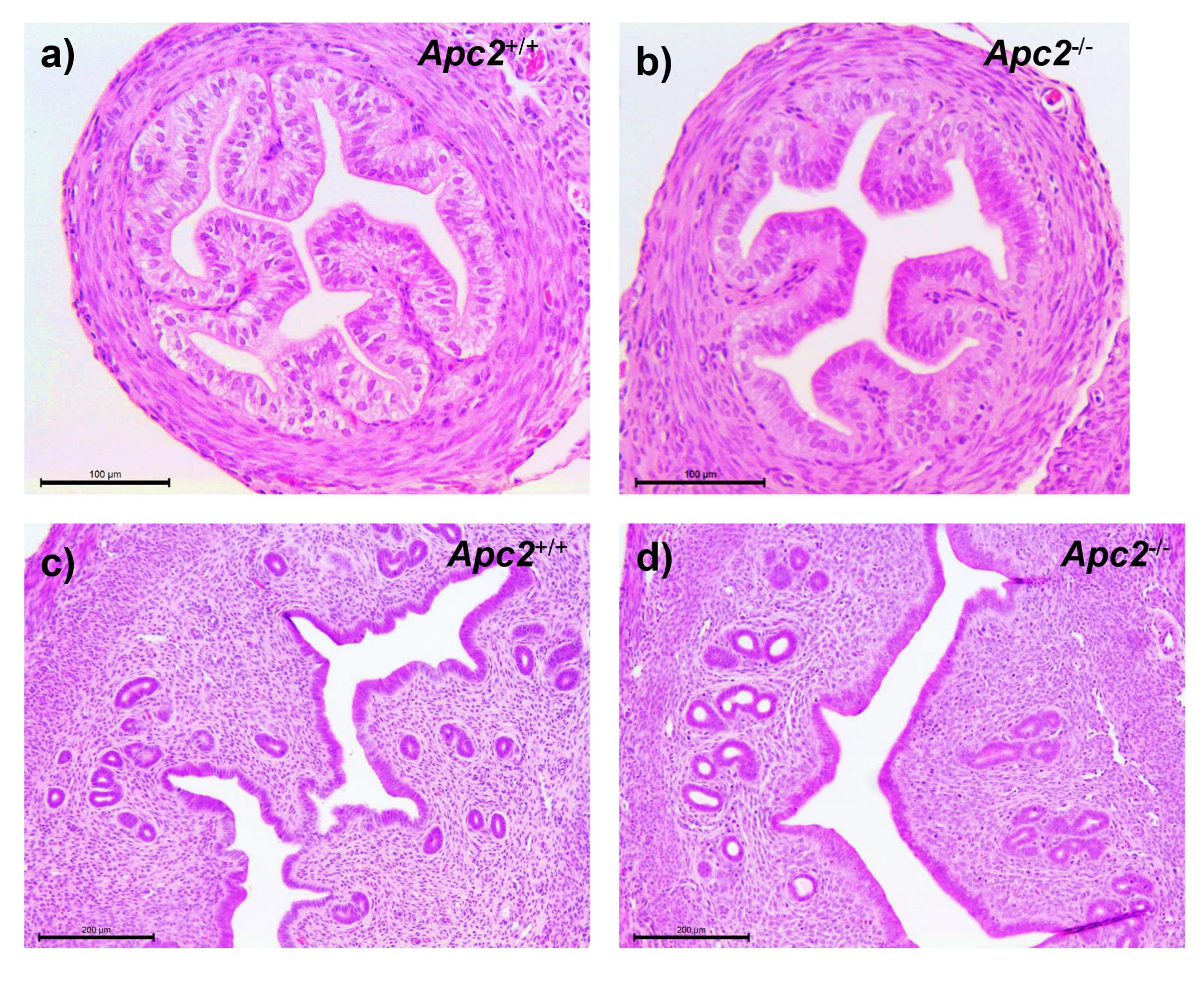

### Supplementary Figure 2

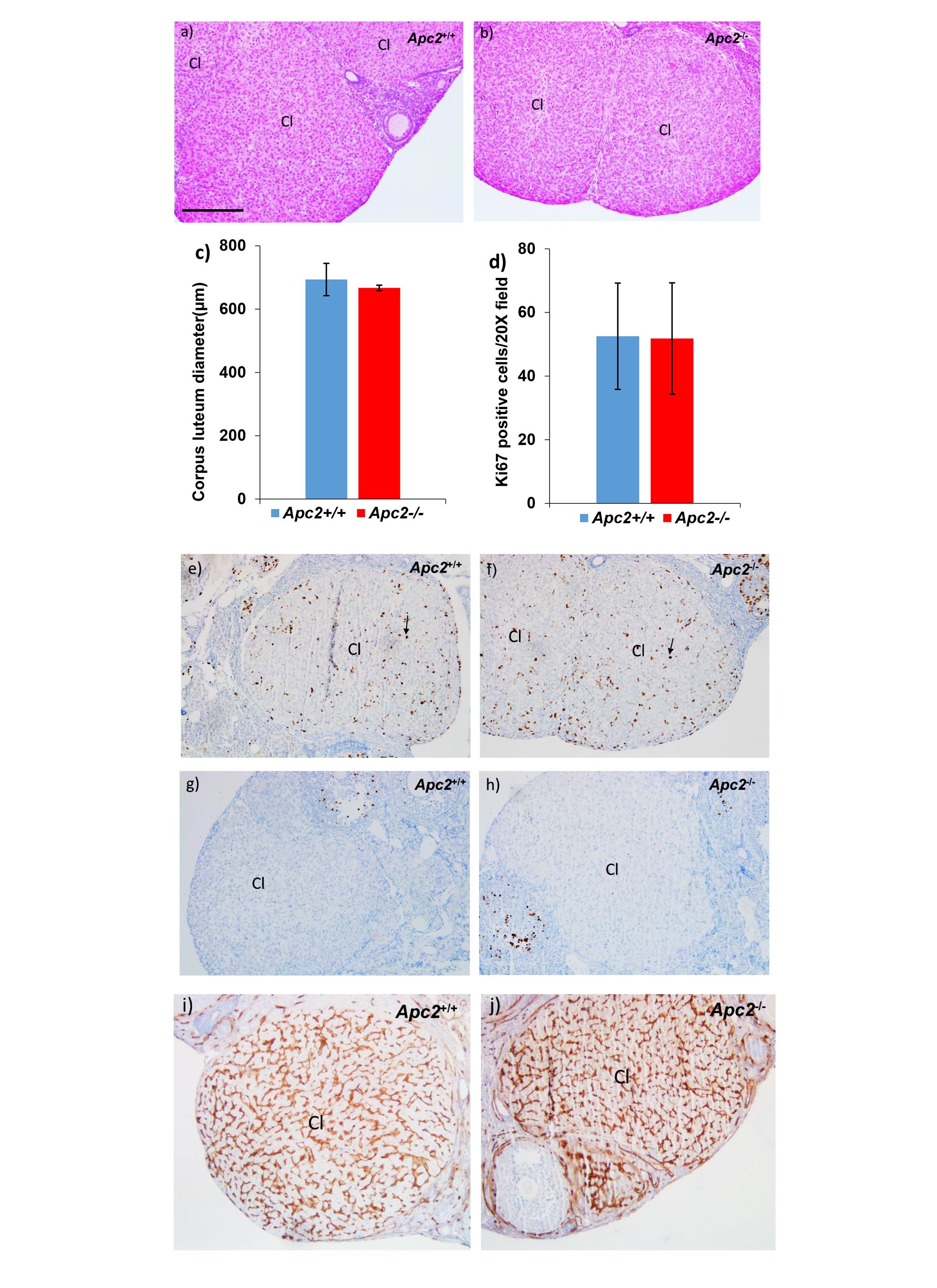

### Supplementary Figure 3

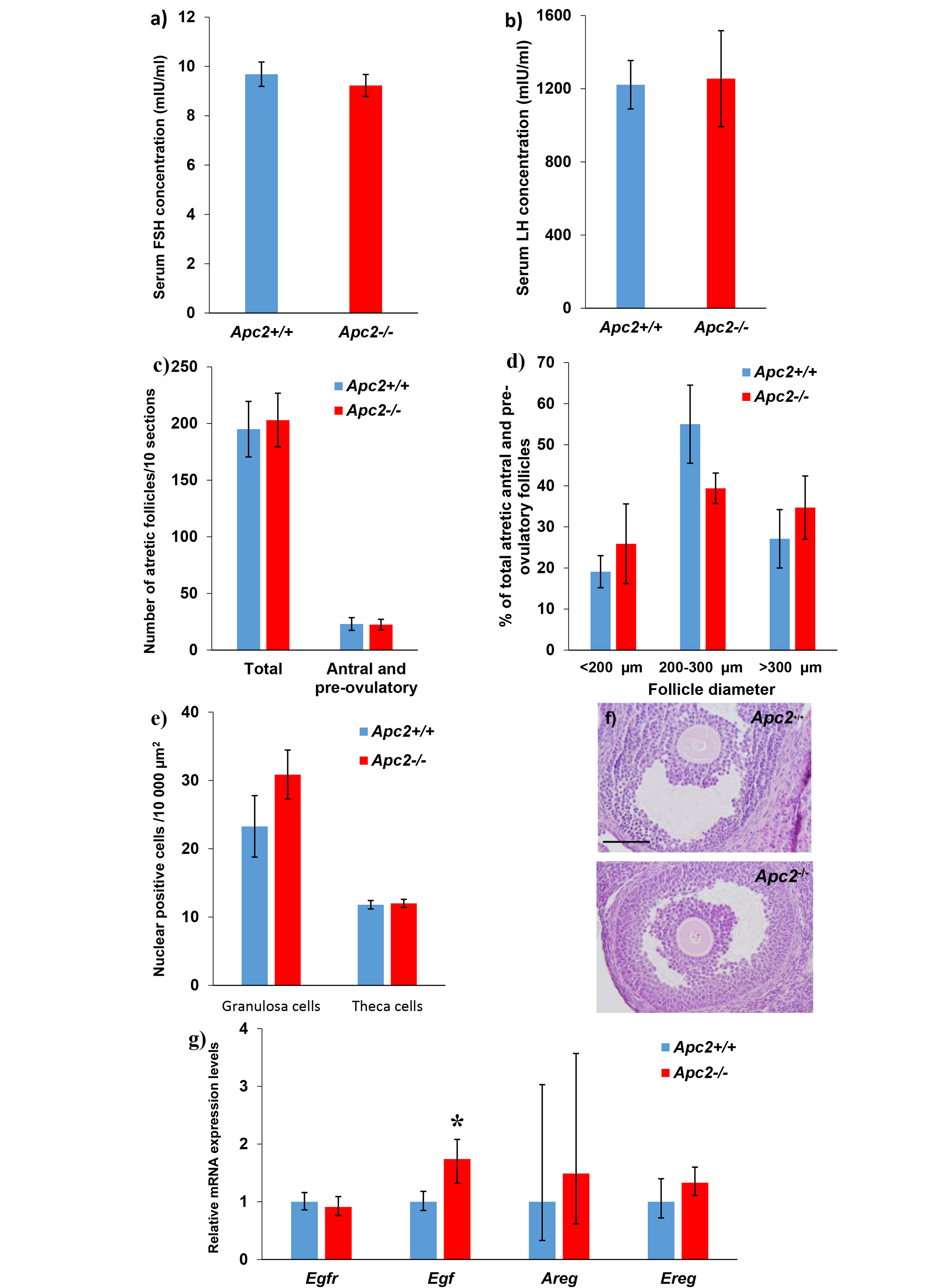

### Supplementary Figure 4

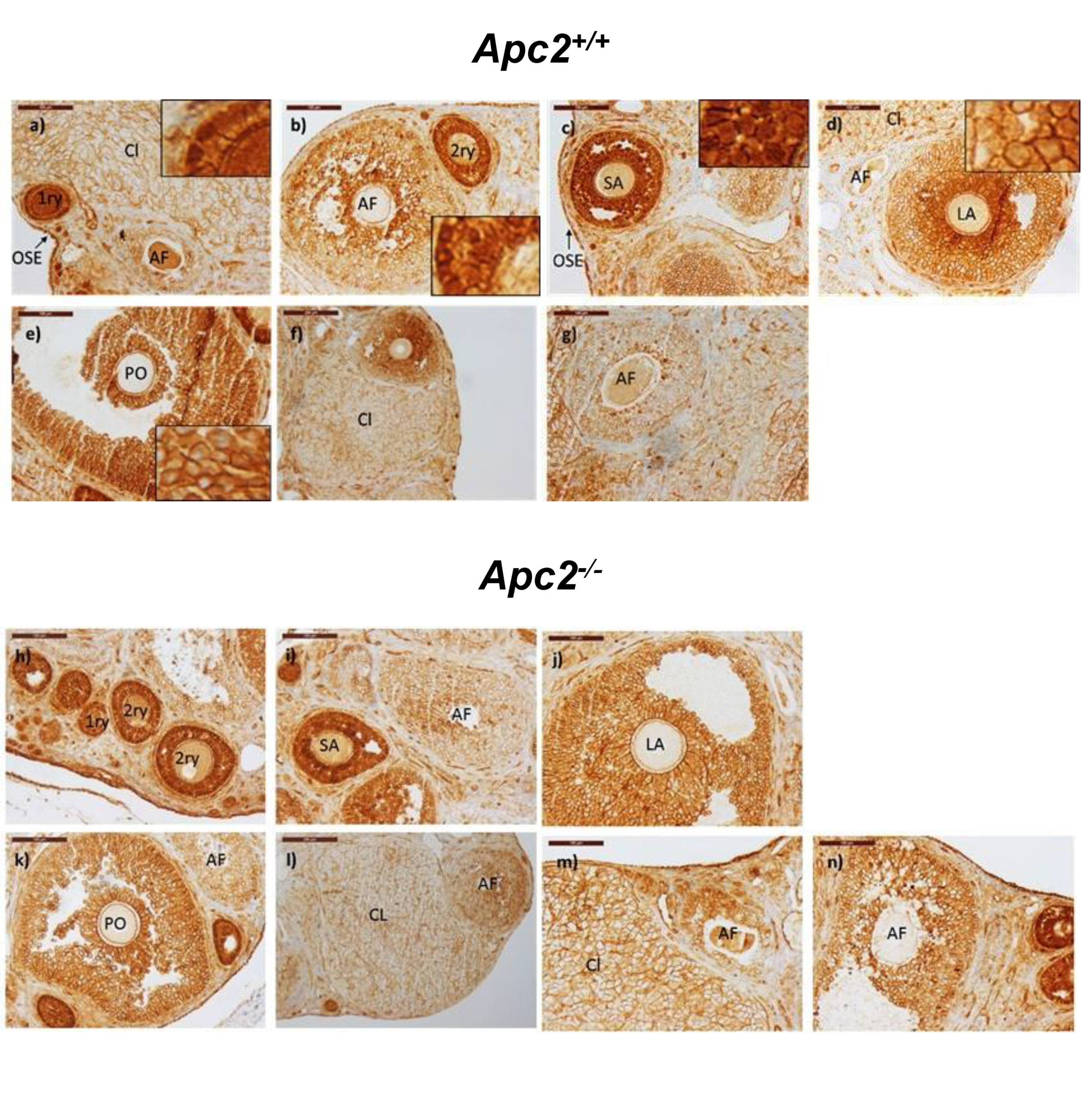

### Supplementary Figure 5

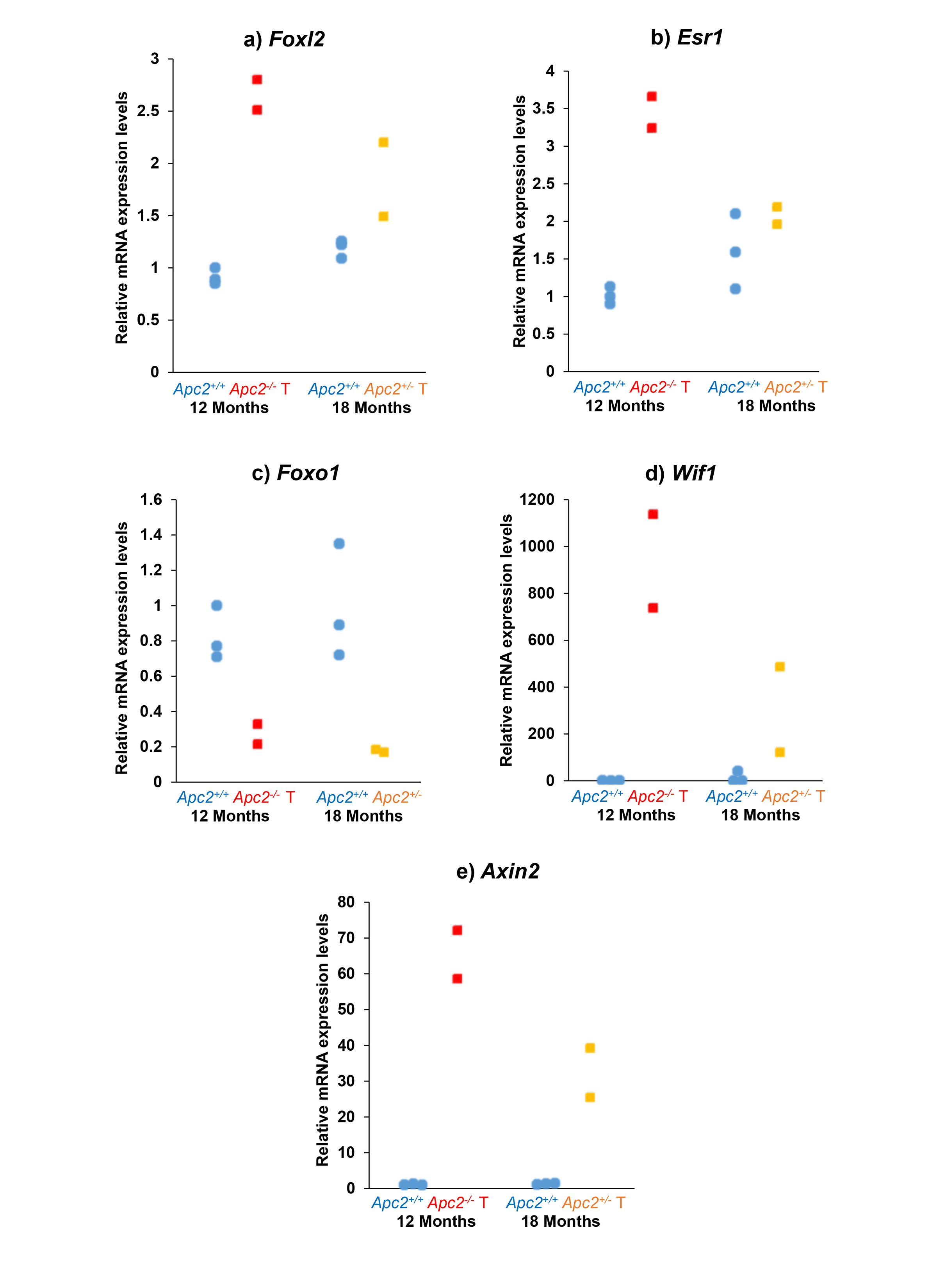
