## Supplementary Tables for "APC2 is Critical for Ovarian WNT Signalling Control, Fertility and Tumour Suppression"

**Supplementary Table 1:** Primer sequences and reaction conditions used in genotyping

| Gene | Primer sequence | Taq polymerase | Buffer | PCR cycle |
| --- | --- | --- | --- | --- |
| *Apc* | Forward primer (5’-3’)  GTTCTGTATCATGGAAAGATAGGTGGTC  Reverse primer (5’-3’)  CACTCAAAACGCTTTTGAGGGTTGATTC | Dream Taq® (ThermoFisher Scientific) | Green | Initial denaturation at 95ºC for 3 minutes followed by 30 cycles (95ºC for 30 seconds, 60ºC for 30 seconds and 72ºC for 1 minute) and a final extension at 72ºC for 5 minutes. |
| *Apc2* | Forward primer (5’-3’)  Wild type primer:  CTCCAAACACAAGATGATCG  Knockout primer:  AGGTCTGAAGAGGAGTTTAC  Reverse primer (5’-3’)  AGCTGTGTCTGATGAGGTG | Go Taq® (Promega) | Clear | Initial denaturation at 95ºC for 3 minutes followed by 30 cycles (95ºC for 30 seconds, 60ºC for 30 seconds and 72ºC for 1 minute) and a final extension at 72ºC for 5 minutes. |

**Supplementary Table 2:** Primary antibodies for immunohistochemistry

| **Primary antibody** | **IgG source** | **Dilution used** | **Blocking serum** | **Secondary antibody and dilution used** |
| --- | --- | --- | --- | --- |
| Β-catenin  Pharmingen  562505 | Mouse | 1/200 | 10% rabbit serum | EnVision+ System- HRP  Labelled Polymer  Anti-mouse |
| Ki67  Abcam  Ab1667 | Rabbit polyclonal | 1/100 | 10% goat serum | Biotinylated goat anti-rabbit immunoglobulins (1/200) |
| Active caspase-3  Cell signalling  9661 | Rabbit polyclonal | 1/200 | 10% goat serum | Biotinylated Goat Anti-Rabbit Immunoglobulins  (1/200) |
| CD34  Abcam  Ab81289 | Rabbit polyclonal | 1/100 | 10% goat serum | Biotinylated Goat Anti-Rabbit Immunoglobulins (1/200) |
| E-cadherin  BD Transduction  610182 | Mouse monoclonal | 1/200 | MOM blocking reagent | EnVision+ System- HRP  Labelled Polymer  Anti-mouse |
| Cytokeratin 18 (CK18) multiepitope cocktail  Progen biotechnik  651134 | Mouse monoclonal | 1/5 | MOM blocking reagent | MOM biotinylated anti-mouse Ig reagent (1/250) |
| PTEN  Cell signalling  9559 | Rabbit monoclonal | 1/200 | 10% goat serum | Biotinylated Goat Anti-Rabbit Immunoglobulin (1/200) |
| p-AKT (Ser 473 D9E)  Cell signalling  4060S | Rabbit Monoclonal | 1/75 | 10% goat serum | Biotinylated Goat Anti-Rabbit Immunoglobulin (1/200) |
| p-FOXO1/3/4  Cell signalling  2599 | Rabbit monoclonal | 1/50 | 10% goat serum | Biotinylated Goat Anti-Rabbit Immunoglobulin (1/200) |
| FOXL2  Abcam  Ab5096 | Goat polyclonal | 1/750 | 10% rabbit serum | Biotinylated rabbit anti-goat immunoglobulin (1/200) |
| ERα | Mouse | 1/500 | MOM blocking reagent | MOM biotinylated anti-mouse Ig reagent (1/250) |
| FOXO1  Cell signalling  14952 | Mouse | 1/500 | 10% goat serum | EnVision+ System- HRP  Labelled anti-mouse |
| Inhibin α (T-17)  Santa cruz  22048 | Goat polyclonal | 1/20 | 10% rabbit serum | Biotinylated rabbit anti-goat immunoglobulin (1/200) |

**Supplementary Table 3:** Taqman® assays used for relative gene expression analysis

| Taqman Gene Expression Assay | Assay ID | Taqman Gene Expression Assay | Assay ID |
| --- | --- | --- | --- |
| *Actb* | Mm00607939_s1 | *Esr1* | Mm00433149_m1 |
| *Apc* | Mm00545877_m1 | *Esr2* | Mm00599819_m1 |
| *Ar* | Mm00442688_m1 | *Fasl* | Mm00438864_m1 |
| *Areg* | Mm00437583_m1 | *Fgf1* | Mm00438906_m1 |
| *Axin2* | Mm00443610_m1 | *Foxl2* | Mm00843544_s1 |
| *Bcl2l11* | Mm00437796_m1 | *Foxo1* | Mm00490671_m1 |
| *Bcl6* | Mm00477633_m1 | *Foxo3* | Mm01185722_m1 |
| *Cd44* | Mm01277163_m1 | *Fshr* | Mm00442819_m1 |
| *Cdkn1b* | Mm00438168_m1 | *Lef1* | Mm00550265_m1 |
| *Ctnnb1* | Mm00483039_m1 | *Lgr5* | Mm00438890_m1 |
| *Cyp11a1* | Mm00490735_m1 | *Lhcgr* | Mm00442931_m1 |
| *Cyp17a1* | Mm00484040_m1 | *Pgr* | Mm00435625_m1 |
| *Cyp19a1* | Mm00484049_m1 | *Tnfsf10* | Mm01283606_m1 |
| *Egf* | Mm00438696_m1 | *Vegfa* | Mm01281448_g1 |
| *Egfr* | Mm00433023_m1 | *Wif1* | Mm00442355_m1 |
| *Ereg* | Mm00514794_m1 |  |  |
